# Evidence for a system in the auditory periphery that may contribute to linking sounds and images in space

**DOI:** 10.1101/2020.07.19.210864

**Authors:** David LK Murphy, Cynthia D King, Stephanie N Lovich, Rachel E Landrum, Christopher A Shera, Jennifer M Groh

## Abstract

Eye movements alter the relationship between the visual and auditory spatial scenes. Signals related to eye movements affect the brain’s auditory pathways from the ear through auditory cortex and beyond, but how these signals might contribute to computing the locations of sounds with respect to the visual scene is poorly understood. Here, we evaluated the information contained in the signals observed at the earliest processing stage, eye movement-related eardrum oscillations (EMREOs). We report that human EMREOs carry information about both horizontal and vertical eye displacement as well as initial/final eye position. We conclude that all of the information necessary to contribute to a suitable coordinate transformation of auditory spatial cues into a common reference frame with visual information is present in this signal. We hypothesize that the underlying mechanism causing EMREOs could impose a transfer function on any incoming sound signal, which could permit subsequent processing stages to compute the positions of sounds in relation to the visual scene.

## Introduction

Every time we move our eyes, our retinae move in relation to our ears. These movements shift the relationship of the visual scene (as detected by the retinal surface) and the auditory scene (as detected based on timing, loudness, and frequency differences in relation to the head and ears). Precise information about the magnitude, direction, starting/ending position, and time of occurrence of each eye movement is therefore needed to connect the brain’s views of the visual and auditory scenes to one another. While there is ample evidence that the brain uses information about eye movements to support visual-auditory integration [1–3], and eye movement signals are known to be rife within the auditory pathway [4–11] as well as association cortex and oculomotor areas [12–23], the computational mechanisms that underlie this process are poorly understood. In particular, it is not well known how the initial *peripheral* stages of sensory signal processing contribute to this process. In hearing (as in other senses), the brain exerts descending control over sensory transduction [24–29]. Thus, the brain has the opportunity to tailor how ascending sensory signals are first encoded, and potentially adjust ascending auditory signals to facilitate linking to the visual system at later stages.

Here, we assessed the potential contributions of one such process. Eye movement-related eardrum oscillations (EMREOs) are a recently discovered phenomenon in which the eardrum oscillates, time-locked to both saccade onset and offset, in the absence of incoming sound [30]. These oscillations likely reflect descending eye movement-related neural commands that influence the ears’ internal motor elements - the middle ear muscles, outer hair cells, or some combination – in a still unknown fashion. In this study, we evaluated the information contained in this signal in human participants. If the mechanism(s) that cause EMREOs are to adjust ascending auditory signals to facilitate localizing sounds in a reference frame shared with vision, these oscillations should vary with both horizontal and vertical dimensions and should depend not only on eye displacements but also at least to a degree on absolute eye positions.

We find that EMREOs consist of a basic waveform time-locked to the onset and offset of eye movements that is modified by the spatial parameters of the eye movements. Displacement of the eyes in the horizontal (i.e. contraversive vs. ipsiversive) dimension has the largest impact, but the vertical dimensions and absolute initial position of the eyes also contribute. The spatial signals contained in EMREOs are sufficiently well-elaborated that target location can be predicted or “read out” from the microphone signals recorded in the ear canal. These findings suggest that the eye-movement information needed to accomplish a coordinate transformation on incoming sounds is fully available in the most peripheral part of the auditory system. While the precise mechanism that creates EMREOs remains unknown, we propose that that underlying mechanism might introduce a transfer function to the sound transduction process, serving to adjust the gain, latency, and/or spectral dependence of responses in the cochlea. In principle, this could allow later stages of processing to extract an eye-centered signal of sound location for registration with the eye-centered visual scene [1].

## Results

We used earbud microphones to record internally-generated sounds in the ear canals of 10 human subjects while they performed eye-movement tasks involving various visual fixation and target configurations, but no external sounds presented (Figure 1). The events of the task are shown in time in Figure 1a. At the beginning of each trial, the subject made a saccade (rapid eye movement) to a visual fixation target (labeled fixation). After a period of 750ms of fixation presentation, a second target appeared and the initial fixation target was turned off. Subjects then made a saccade to that second target and maintained fixation for another 200 ms (labeled target). If subjects did not maintain pre- and post-saccade fixation for at least 200ms, the trial was discarded. The two periods of fixation ensured that the eyes were stable for a period before and after the fixation-to-target saccade so baseline ear-canal recording measures could be established without eye movement. We analyzed the ear-canal recordings associated with this second saccade.

**Figure 1.**
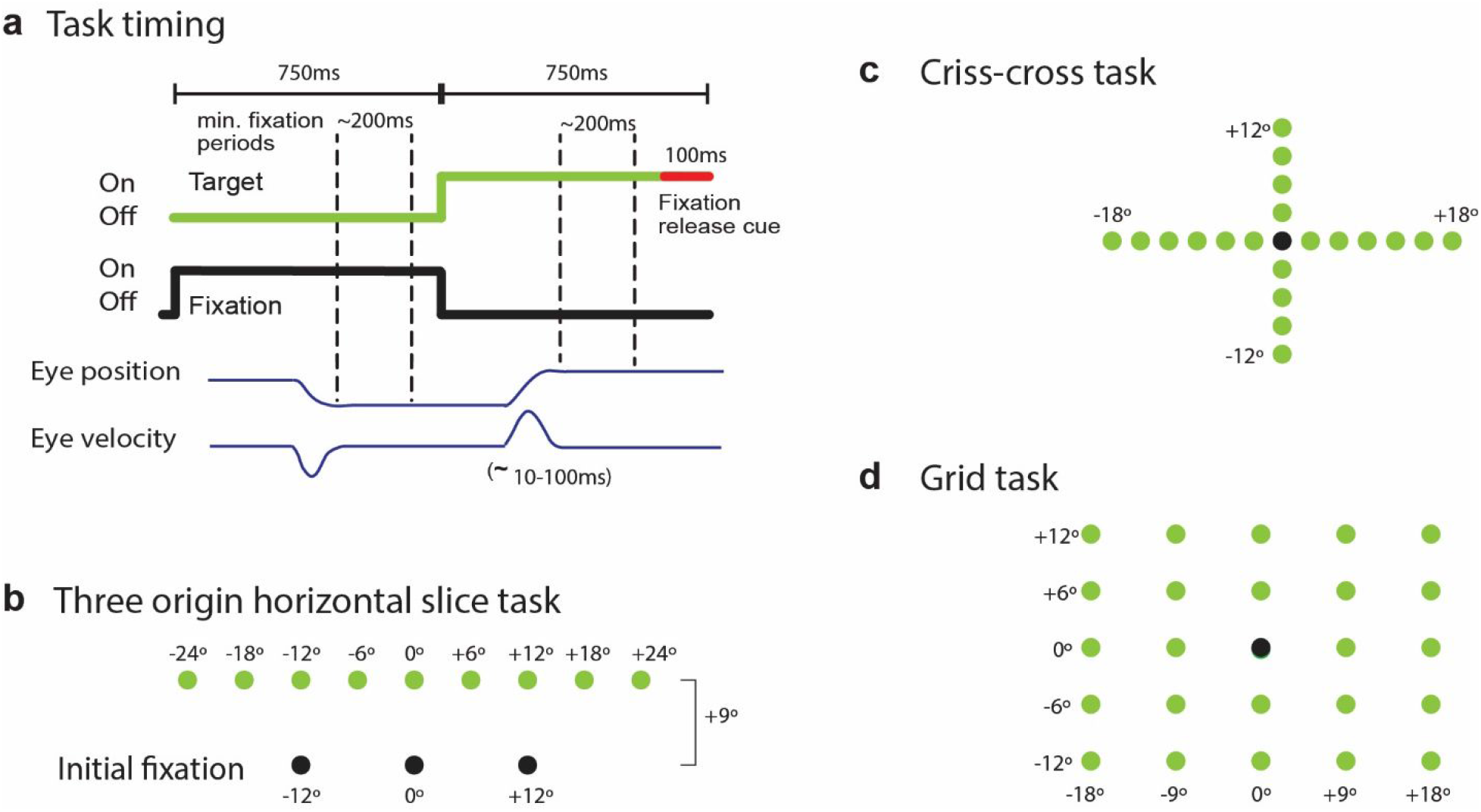
Events of the tasks in time and space. a. Task events across time. Each trial began with the onset of an initial fixation cue (labeled fixation). Participants made saccades to the fixation point, then maintained fixation for a minimum of 200 ms. The fixation cue was turned off and a new target requiring a second saccade was turned on. Participants then fixated on the second target (labeled target) for another 200 ms, at which point the target turned red indicating that the trial was over. The ear-canal recordings were analyzed in conjunction with the second saccade. b-d. Spatial layouts of fixations (black) and targets (green) for the three task designs used in this study.

Three different task designs are shown in panels b-c, and were chosen to focus on three questions: 1) what are the respective contributions of relative eye displacement vs absolute initial/final position of the eyes to the EMREO signal (three-origin task)? 2) what are the respective contributions of purely horizontal vs purely vertical saccade displacements to EMREOs (criss-cross task)? And 3) how do the horizontal and vertical eye displacement components of EMREOs interact for saccades spanning full 2-dimensional space, i.e., oblique saccades (grid task)? The answers to these questions provide a sense of the full spatial dependence of these eardrum oscillations.

Using the three-origin horizontal slice task, we first replicated our original findings by analyzing only the trials that used similar targets as previously published work [30]. These consisted of a horizontal target array at a single azimuth or horizontal “slice” with a fixed vertical offset from a single center fixation position. Then, we extended our original research by including additional fixation positions as well as additional targets in the analysis. Results for the conditions that most closely matched our previous study are shown for an example subject’s left ear in Figure 2a-f. Panels a and b show average horizontal eye position as a function of time for each of the targets between −24 and 24 degrees (the original study used −18 to +18 degrees, as well as a 6 degree vertical offset compared to 9 degrees here). The traces are color coded by the target location, with the inset showing the average eye traces in two dimensions. In panel a, the saccades are aligned on saccade onset, defined as the first peak in the third derivative of eye position over time (eye jerk), or the point in time when the change in eye acceleration is highest. In panel b, the saccades are aligned on saccade offset, defined as the second peak in eye jerk, or when the change in eye deceleration is greatest. Panels c and d show the average velocity traces associated with these saccades. Finally, panels e and f are the average microphone signals recorded in the left ear at the time these saccades were made, color-coded by visual target.

**Figure 2.**
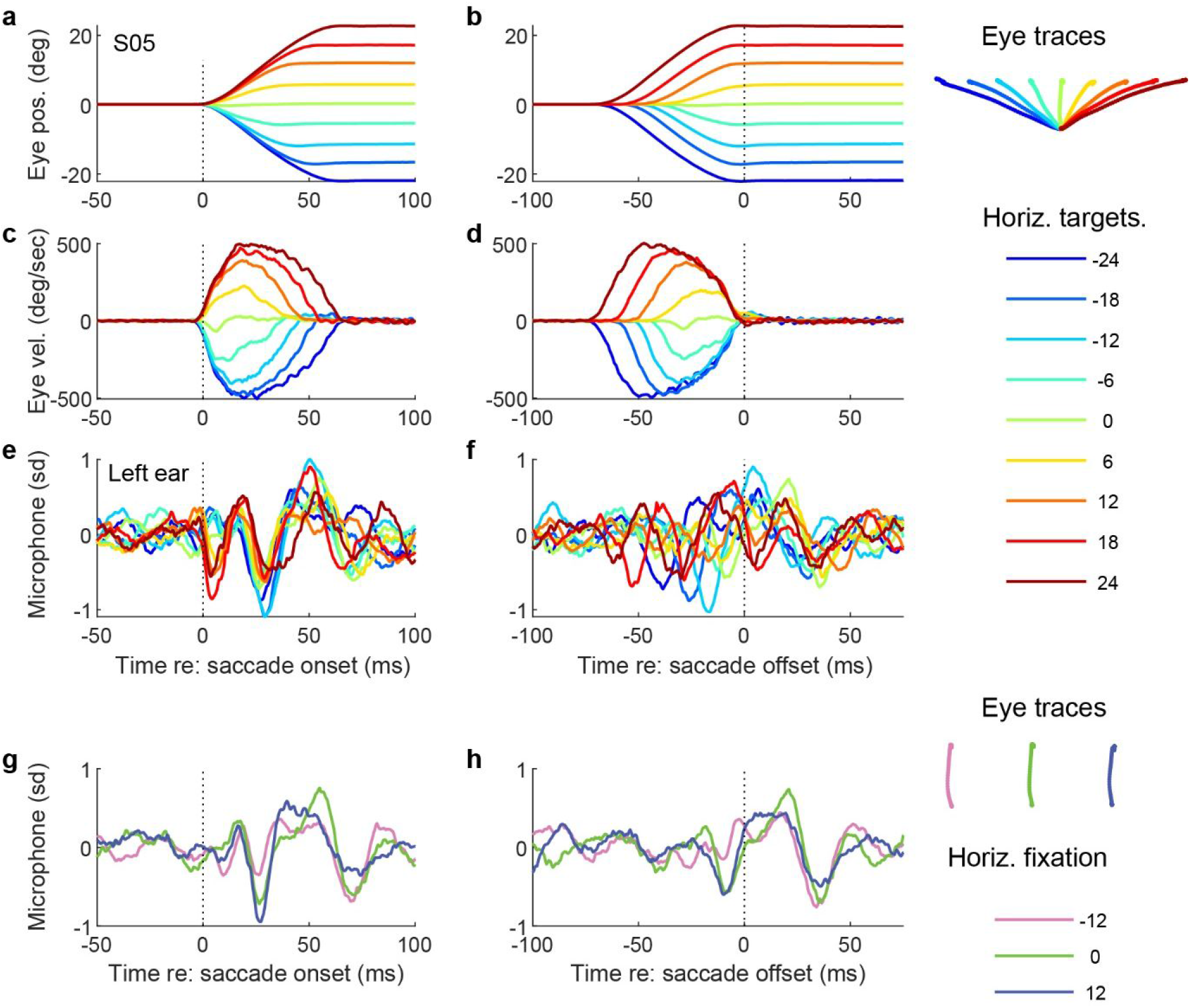
Results for one participant’s left ear for the three origin horizontal slice task. The left column shows results aligned on saccade onset. The right column shows the results aligned on saccade offset. Color corresponds to various target and fixation combinations, as illustrated by the inset eye traces. Panels a-f involve only the center initial fixation position and its secondary targets (replicating previous research). The different shades of blue/green are ipsiversive eye movements and the red/yellow are contraversive, regardless which ear is depicted. Panels g-h show results of microphone recordings from all three initial fixation positions (12, 0, −12 degrees) and their corresponding straight vertical saccades. All microphone data was z-scored relative to the mean and standard deviation of a time window −150 to −120 ms prior to saccade onset, i.e. during a period of steady fixation. Units on the y axis are therefore in units of standard deviation relative to this baseline.

As described in our previous work, an internally-generated oscillation (i.e. sound) occurs in conjunction with each eye movement. The phase and amplitude vary with the direction and magnitude of the saccade, and there is evident of phase-resetting at both saccade onset and offset. The phase resetting at saccade offset is particularly apparent in panel f, where the first peaks or troughs roughly line up with each other after offset, whereas beforehand they are staggered according to the associated target.

The current study builds on our original findings by considering the contributions of absolute initial/final eye position to the EMREO pattern. We evaluated the results for all three fixation locations, the straight ahead fixation as well as 12 degrees on either side, all in the 0 degree vertical plane. To assess the impact of initial/final eye position on the EMREO signal, we compared ear-canal recordings from saccades that had the same direction and amplitude, but different starting and ending positions. Panels 2g and 2h show an example of average microphone traces for three such fixation/target combinations. Overall, the shape of the microphone trace appears similar but not identical for the same eye displacement from different absolute positions.

To statistically quantify the pattern of dependence on eye position and displacement, we deployed a regression analysis in which the microphone signal at each moment in time Mic(t) was fit to the horizontal initial eye position (H) and change in eye position (ΔH) on that trial as follows:

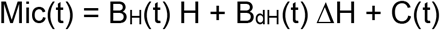

with B_H_ and B_dH_ expressing the contributions related to initial horizontal eye position and eye displacement respectively, and C(t) providing a time-varying constant term that can be thought of here as the average oscillation regardless of horizontal saccade metrics (see Methods, equation 1). To eliminate a correlation between H and ΔH, we included only trials for which the horizontal target displacements occurred for all three initial fixation positions, i.e. −12° to +12° targets for the 0° fixation position, −24° to 0° targets for the −12° fixation, and 0° to 24° targets for the +12° fixation position.

Figure 3 shows the regression terms across time (aligned to saccade onset) for the 10 subjects in our study. Red lines show the data for the right ear and blue for the simultaneously recorded left ear. Thickened portions indicate periods of time when the 95% confidence intervals for the given regression coefficient did not include zero.

**Figure 3.**
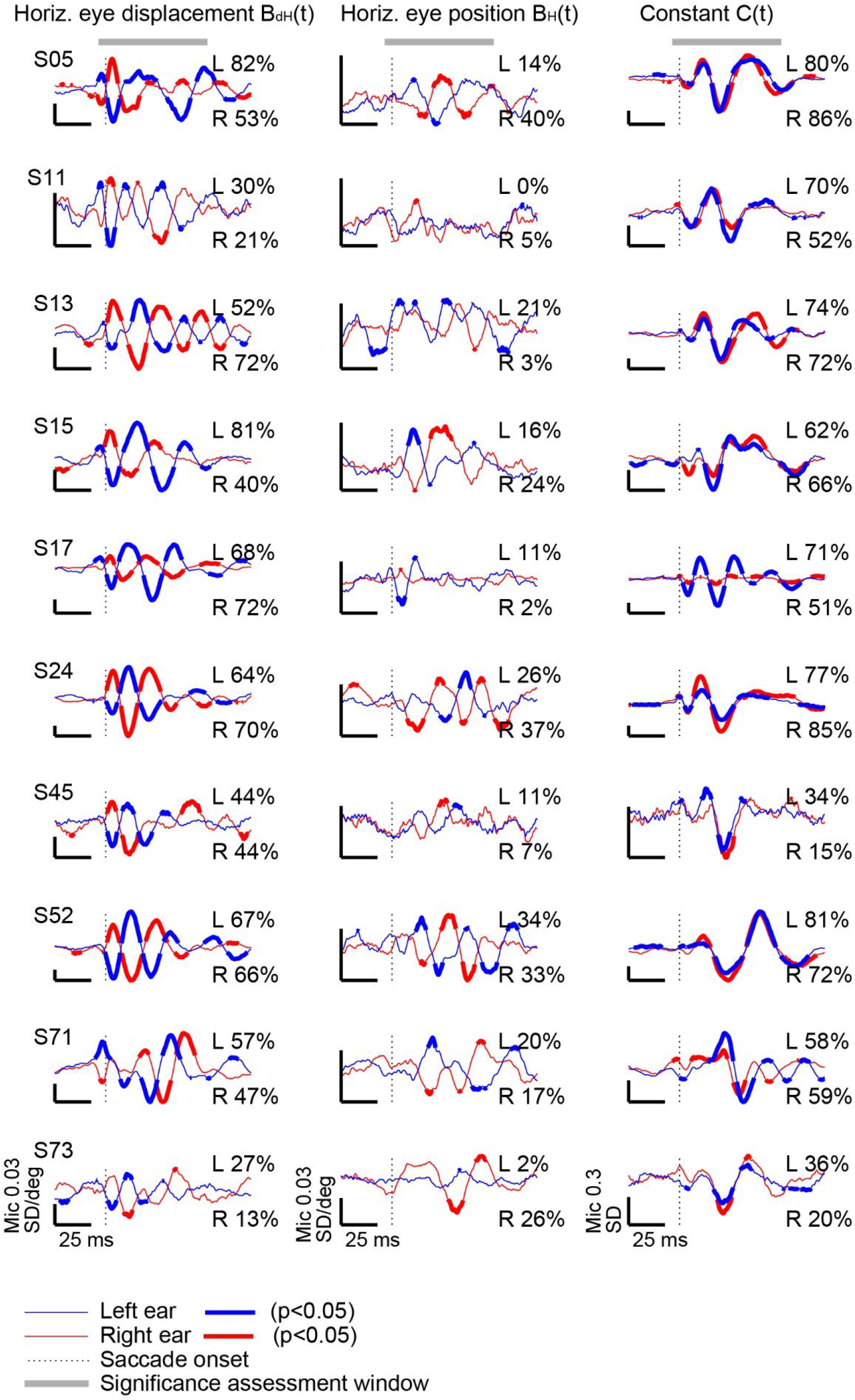
Regression results for all participants for the three origin horizontal slice task. Thick line segments indicate epochs of time in which the corresponding regression term differed from zero (p<0.05). Percent values reflect the percentage of time points in the window −5 ms to 70 ms with respect to saccade onset (the significance assessment window) that differed from zero with 95% confidence. Y axis units are standard deviations per degree (horizontal eye displacement and horizontal initial eye position columns) or simply standard deviations (“constant” column) and vary across panels as indicated by the calibration bars.

The left column shows the regression coefficients associated with horizontal eye displacement (BH(t)). These traces show a characteristic pattern that is present in all subjects: a peak-trough-peak oscillation or trough-peak-trough oscillation depending which ear is being recorded. The phase is inverted between ears, indicating that the relevant aspect of eye displacement is whether it is ipsiversive or contraversive to the ear canal where the signal is recorded. Every individual participant shows this characteristic pattern. Other than modest differences in overall amplitude (note the different calibration bars for each subplot), subject variability occurs mainly in the duration and latency of the overall pattern. For example, the overall duration of the oscillation is the longest in S13, lasting ~100 ms. Most subjects show a clear peak/trough just prior to saccade onset (e.g. S05 and S71), while some do not (e.g. S24).

The right column shows the constant term of the regression equation (*C(t)*), reflecting the average oscillation that occurs regardless of the metrics of the associated eye movement in the horizontal dimension. Here, the left and right ear traces are extremely similar to each other in most individual subjects (except S17, who has a magnitude difference across the two ears). Subjects differ from one another in the timing/duration of this element of the effect. For example, S52 exhibits a fairly late oscillation in the value of the constant term, beginning about 25 ms after saccade onset. However, there is considerable consistency across subjects, with most showing a trough-peak-trough pattern starting at saccade onset. (Note: All eye movements in this task involved a vertical component, but that vertical component did not vary, so any dependence of the EMREO on the vertical dimension would be captured by the constant term in this version of our regression analysis. We will return to this issue below using the criss-cross and grid tasks to provide a resolution.)

The middle column shows the corresponding data for the horizontal initial/final eye position dependence. The microphone signal’s dependence on starting/ending eye position was smaller and more variable across subjects than either of the other two regression components. When present, it exhibited a counter-phase pattern across the two ears similar to that seen for the horizontal displacement dependence.

Quantifying the statistical significance at the individual subject level is non-trivial because the microphone signal at a given time point is not independent of the signal at adjacent time points. To provide a general sense of significance, we computed the percentage of individual time points for which the relevant regression terms differed from zero at the 95% confidence intervals during a time period of 5 ms before to 70 ms after saccade onset, dubbed the significance assessment window, as indicated by the horizontal bars in the top row of graphs. This range was chosen to roughly capture the full oscillations of all subjects. Note that this percentage cannot reach 100% because microphone signals oscillate around zero and thus the 95% confidence intervals will include zero during those times. However, it provides a useful benchmark. The percent-significant-time points for each individual subject’s left and right ear traces are shown on each individual panel. For the constant terms, these percentages range from 15-86%, and for the horizontal displacement terms, they range from 13-82%. Using a decision criterion that each individual subject must show at least 10% significant points for at least one ear, all 10 subjects easily surpassed this and can be said to exhibit a significant dependence of the microphone signal on horizontal eye displacement as well as showing a significant constant term.

The pattern is weaker for the horizontal initial/final position component: 9 of 10 reached the criterion (except S11), but it should be noted that two do so just barely (S17, S45). We conclude that on average there does exist a dependence on horizontal initial/final position but that this dependence is weaker, more variable, and potentially less regular than the dependence on horizontal eye displacement. The irregular nature of the impact of this factor suggests that it may require methods other than regression to fully characterize.

We next explored eye displacement sensitivity in two dimensions. In the criss-cross task (Figure 1c), we assessed purely horizontal and purely vertical saccades initiated from a single central fixation point. Figure 4 shows the results from the same example subject shown in Figure 2. The top four panels (4a-d) show the average horizontal eye position and velocity traces for a slice of horizontal targets with no vertical offset and with a finer target spacing of 3 degrees, aligned on saccade onset (left panels) and offset (right panels). The corresponding microphone signals are shown in panels 4e and f. The results are reasonably similar to our original findings, which used a non-zero vertical offset (of the target from the initial eye position), confirming that the previously observed horizontal displacement dependence applies even in the absence of a vertical component to the saccade. The bottom 6 panels (4g-l) show the results for purely vertical displacements to targets. Again, there is a clear oscillation associated with vertical saccades, with this subject showing a larger signal for upward than for downward saccades. The oscillation continues for approximately 50 ms or more after the saccade has ended.

**Figure 4.**
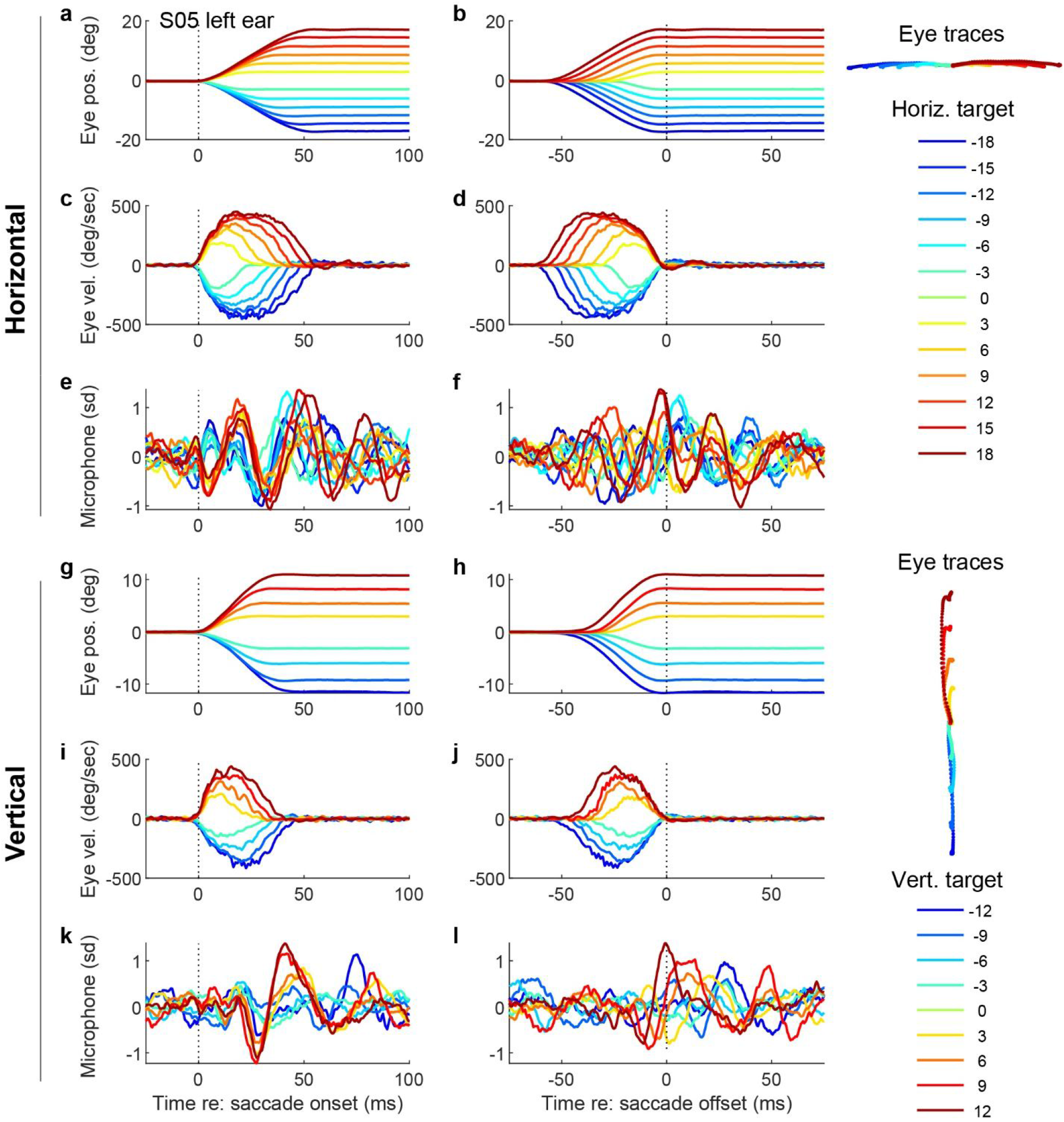
Results from one participant’s left ear (same participant as in Figure 2) in the criss-cross task. Conventions similar to Figure 2. Panels a-f are results for purely horizontal saccades and panels g-l are results for purely vertical saccades.

To fit this data statistically, we again deployed a regression model, this time omitting the term for the horizontal initial/final position and instead incorporating a term for the vertical displacement:

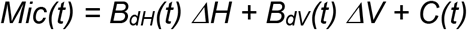

The trials with horizontal targets and the trials with vertical targets were included in the same regression fit as they were collected in the same sessions. Figure 5 shows the results for all subjects, in a similar format as Figure 3.

**Figure 5.**
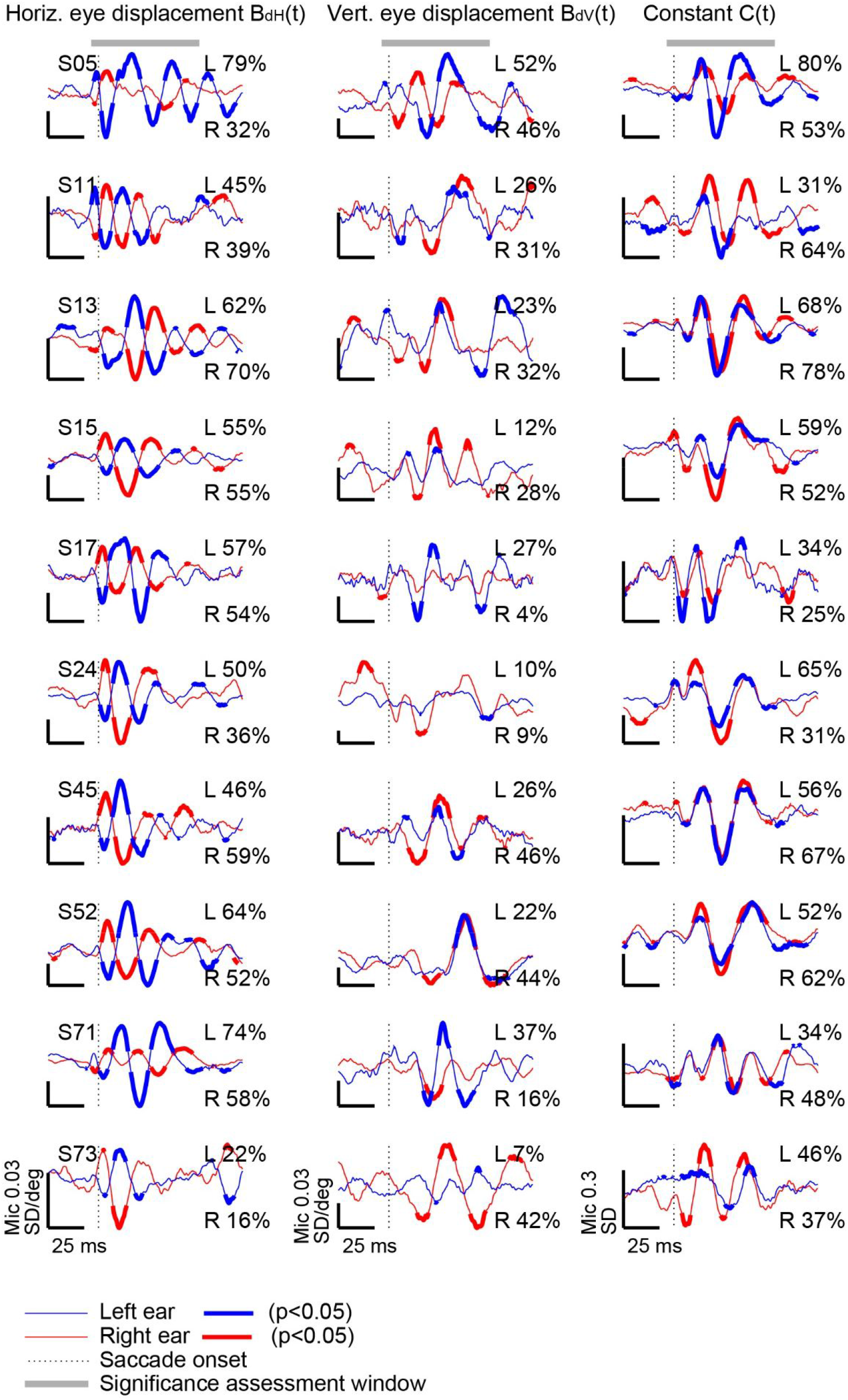
Regression results for all participants for the criss-cross task. Conventions similar to Figure 3.

The dependence of the microphone signal on the horizontal displacement for purely horizontal saccades (left column) is quite similar to those seen in our original study [30] as well as to the three-origin horizontal slice task results presented in Figure 3. Likewise, the constant component (right column) is also similar overall. All 10 subjects meet criteria for showing a significant overall effect for each of these factors.

The middle column shows the dependence on purely vertical saccade displacement. Using our earlier established criterion (i.e. at least one ear exhibiting significance for this factor 10% of the time), a vertical displacement signal is present in all 10 subjects. However, the pattern of effects for vertical saccade amplitude was substantially more variable across subjects and usually occurred much later than the effects for horizontal saccade amplitude (see for example S52). Note that the sensitivity to vertical saccade amplitude was in phase between ears in nearly all subjects, in contrast to the out-of-phase sensitivity to horizontal saccade amplitude between ears.

Next we considered the question of how these distinct dependencies on horizontal and vertical saccade amplitude interact with each other. How do the EMREOs vary in full two-dimensional space? Can the microphone signal for a full range of oblique saccades be predicted from the individual horizontal and vertical eye movement displacement components? To evaluate this, we tested subjects on one more task, a 2D grid of targets which incorporated combined horizontal and vertical displacements from a single fixation origin (see Figure 1d). As shown in Figure 6, the regression coefficient curves for horizontal displacement (*B_dH_(t)*) and the constant terms (*C(t)*) were very similar in the two tasks (grand average across the population, black vs. green traces). The vertical displacement term (*B_dv_(t)*) was more variable but still quite similar across the two tasks.

**Figure 6.**
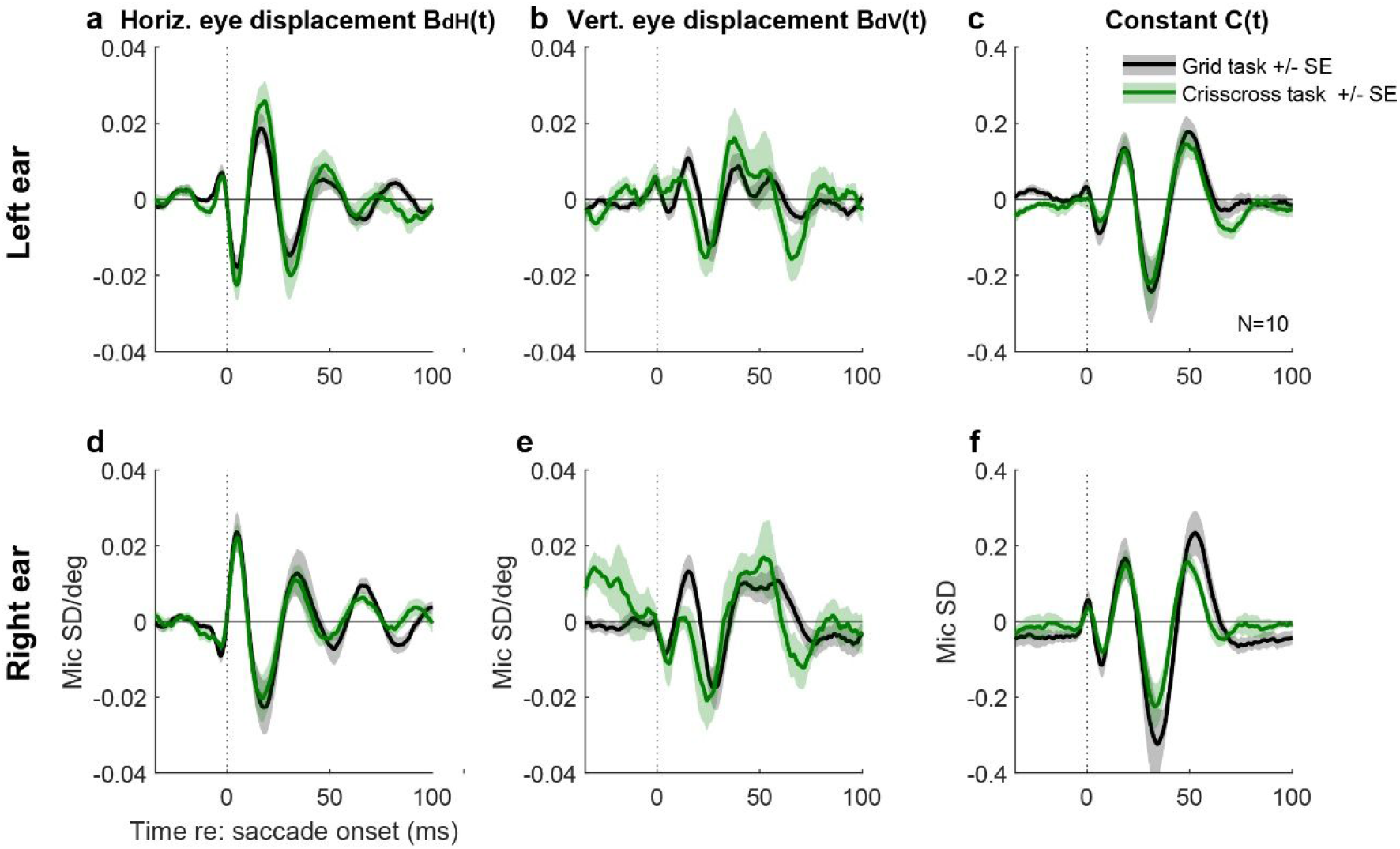
Grand average regression coefficient curves for the grid and criss-cross tasks correspond well. Black and green lines represent the average coefficient values across 10 subjects; shading reflects the standard error across subjects.

Given the similarity of these results, we surmised that it should be possible to predict the microphone signal observed in the grid task using the data from the criss-cross task. Figure 7 shows the grand average microphone signal associated with each targets in the grid task (black traces). The location of each trace corresponds to the physical location of the associated target in the grid task. The superimposed predicted wave forms (red traces) were generated from the B_dH_(t), B_dv_(t), and C(t) regression coefficients fit to the criss-cross data, then evaluated at each target location and moment in time to produce predicted curves for each of the locations tested in the grid task.

**Figure 7.**
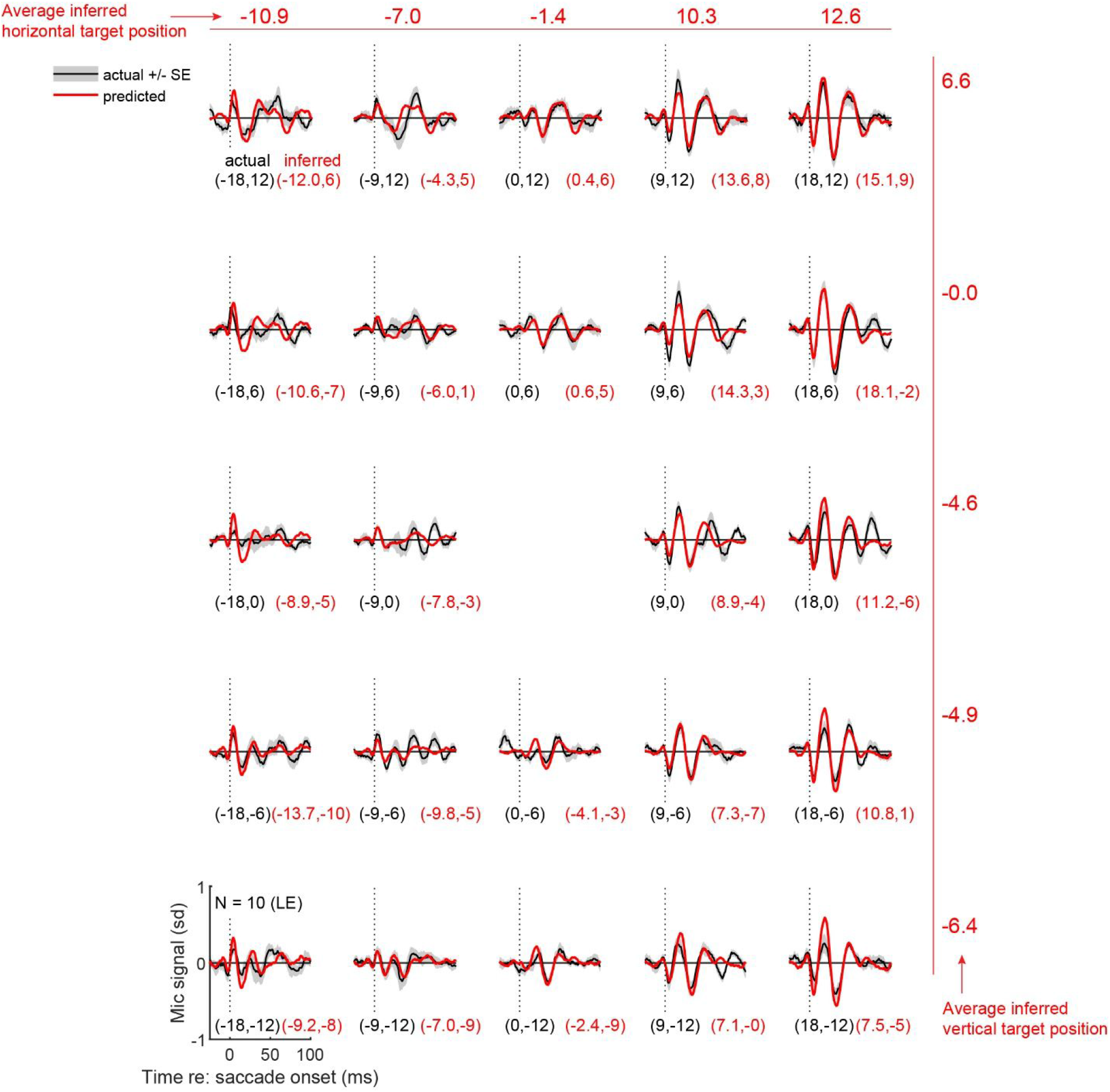
Combined results for all participants’ left ears for the grid task (black lines), with predicted results from the criss-cross task (red lines). The traces indicate the grand average of all the individual participants’ mean microphone signals, with the shading indicating +/- the standard error across participants. Black values in parentheses are the actual horizontal and vertical coordinates for each target in the grid task. Corresponding red values indicate the inferred target location based on solving a multivariate regression which fits the observed grid task microphone signals in the significance assessment window (−5 to 70 ms with respect to saccade onset) to the observed regression weights from the criss-cross task for each target location. The averages of these values in the horizontal and vertical dimensions are shown across the top and right side.

Overall, there is good correspondence between the predicted and observed curves, even for oblique saccade displacements that were not tested in the criss-cross task, i.e. the top left, top right, bottom left, and bottom right corners of this figure. This illustrates two things: a) the EMREO is reproducible across task contexts, and b) the horizontal and vertical displacement dependencies interact in a reasonably linear way, so that the signal observed for a combined horizontal-vertical saccade can be predicted as the sum of the signals observed for straight-horizontal and straight-vertical saccades with the corresponding component amplitudes.

Given that it is possible to predict the microphone signal from one task context to another, it should be possible to infer the target and its associated eye movement from the microphone signal itself. To do this, we again used the weights for the regression equation

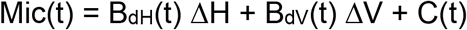

from the criss-cross data. We then used the Mic(t) values observed in the grid task to solve this system of multivariate linear equations across the significance assessment window (time interval −5 to 70 ms with respect to the saccade) to generate the “read out” values of ΔH and ΔV associated with each target’s actual ΔH and ΔV. We conducted this analysis on the left ear and right ear data separately. The left ear results of this analysis are seen in each of the individual panels of Figure 7; the black values (e.g. −18, 12) indicate the actual horizontal and vertical locations of the target, and the associated red values indicate the inferred location of the target. Across the top of the figure, the numbers indicate the average inferred horizontal location, and down the right side, the numbers indicate the average inferred vertical location. These results indicate that, on average, the targets can be read out in the proper order, but the spatial scale is compressed: the average read-out values for the +/-18 degree horizontal targets are +/− ~11-12 degrees, and the averages for the vertical +/− 12 degree targets are +/− ~6-7 degrees.

We wondered whether the accuracy of this read-out could be improved by combining signals recorded in each ear simultaneously – given that it is known that the brain uses binaural computations for reconstructing auditory space. We first considered a binaural difference computation – subtracting the right ear microphone recordings from the left, eliminating the part of the signal that is common between the two ears. Figure 8A illustrates the inferred location of the target (red dots) connected to the actual location of the target (black dots). Generally, the horizontal dimension is well ordered whereas the vertical dimension has considerable shuffling. This can also be seen in panels B and C which show the relationship between the inferred target location and the true target location broken out separately for each dimension. The correlation between inferred and actual target is higher in the horizontal dimension than the vertical dimension. This makes sense because the binaural difference computation serves to diminish the contribution from aspects of the signal that are in phase across the two ears, such as the dependence on vertical eye displacement. Improvement in the vertical readout can be achieved by instead averaging the signals across the two ears (D). A hybrid readout operation in which the horizontal location is computed from the binaural difference, and the vertical location is computed from the binaural average, produces a modest improvement in the overall reconstruction of target location. Overall, these results parallel human sound localization which relies on a binaural difference computation in the horizontal dimension (and is more accurate in that dimension), vs. potentially monaural spectral cues for the vertical dimension (which is less accurate) [31, 32].

**Figure 8.**
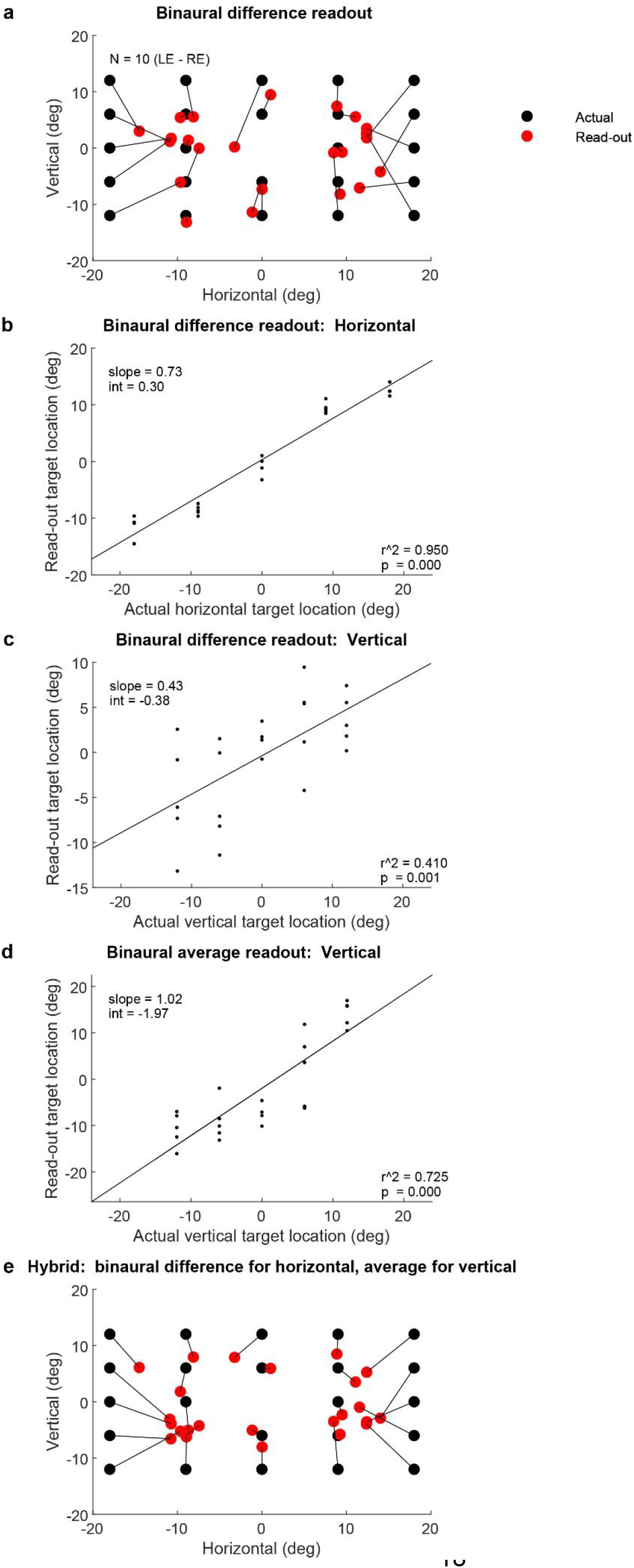
Reading out target location from the grid task microphone signals based on the regression coefficients from the criss-cross task. a. Inferred target location (red) compared to actual target location (black), based on the difference between the two ears’ microphone signals (left ear – right ear). b. Horizontal component of the read-out target vs the actual horizontal component (left ear – right ear microphone signals). c. Same as (b) but for the vertical component. d. Vertical component computed from the average of the two ears. e. A hybrid read-out model using binaural difference in the horizontal dimension and binaural average in the vertical dimension.

## Discussion

Sound locations are inferred from head-centered differences in sound arrival time, loudness, and spectral content, but visual stimulus locations are inferred from eye-centered retinal locations [31, 32]. Information about eye movements with respect to the head/ears is critical for connecting the visual and auditory scenes to one another [1]. This insight has motivated a number of previous neurophysiological studies in various brain areas in monkeys and cats, all of which showed that changes in eye position affected the auditory response properties of at least some neurons (Inferior colliculus: [7–11]; auditory cortex: [4–6]; superior colliculus: [17–23];frontal eye fields: [12, 33]; intraparietal cortex: [13–16]). These findings raised the question of where signals related to eye movements first appear in the auditory processing stream. The discovery of EMREOs [30] introduced the intriguing possibility that the computational process leading to visual-auditory integration might begin in the most peripheral part of the auditory system. Here we show that the signals present in the ear exhibit the properties necessary for playing a role in this process: these signals are influenced by both the horizontal and vertical components of eye movements, and display signatures related to both change-in-eye-position and the absolute position of the eyes in the orbits.

Our present observations raise two key questions: what causes EMREOs and how do those causes impact hearing/auditory processing? The proximate cause of EMREOs is likely to be one or more of the known types of motor elements in the ear: the middle ear muscles (stapedius and tensor tympani), which modulate the motion of the ossicles [34–36], and the outer hair cells, which modulate the motion of the basilar membrane [37]. One or more of these elements may be driven by descending brain signals originating from within the oculomotor pathway and entering the auditory pathway somewhere along the descending stream that ultimately reaches the ear via the 5^th^ (tensor tympani), 7^th^ (stapedius muscle), and/or 8^th^ nerves (outer hair cells) [see refs: 24, 25–29] for reviews). Efforts are currently underway in our laboratory to identify EMREO generators/modulators [38, 39].

Uncovering the underlying mechanism should shed light on a second question: does the temporal pattern of the observed EMREO signal reflect the time course and nature of that underlying mechanism’s impact on auditory processing? It is not clear how an oscillatory signal like the one observed here might contribute to hearing. However, it is also not clear that the underlying effect is, in fact, oscillatory. Microphones can only detect signals with oscillatory energy in the range of sensitivity of the microphone. It is possible that the observed oscillations therefore reflect ringing associated with a change in some mechanical property of the transduction system, and that that change has a non-oscillatory temporal profile (Figure 9A). Of particular interest would be a ramp-to-step profile in which aspects of the middle or inner ear shift from one state to another during the course of a saccade and hold steady at the new state during the subsequent fixation period. This kind of temporal profile would match the time course of the saccade itself.

**Figure 9.**
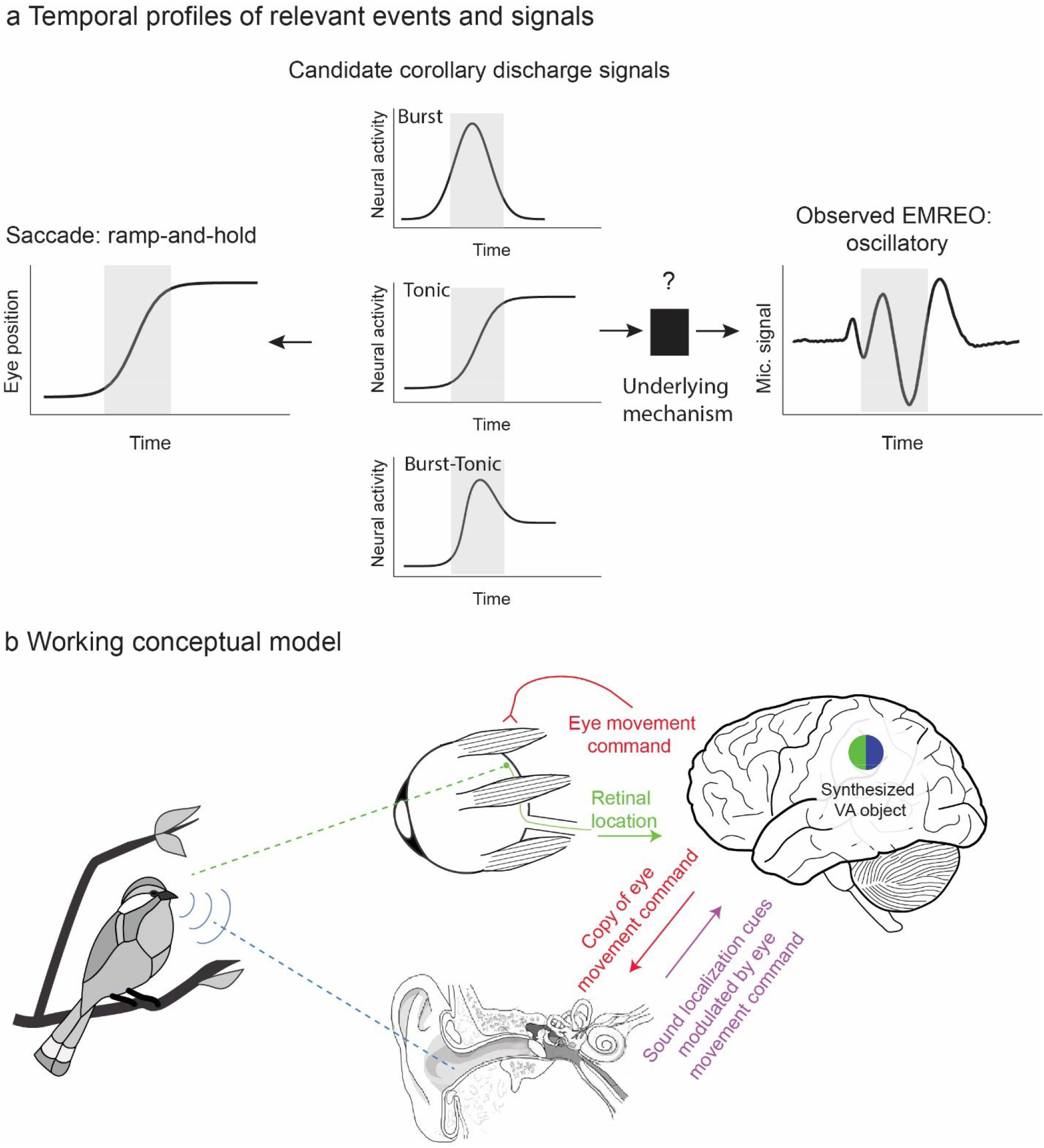
Temporal profiles of relevant signals and working conceptual model for how EMREOs might relate to our ability to link visual and auditory stimuli in space. A. Temporal profiles of signals. The EMREO is oscillatory whereas the eye movement to which it is synchronized involves a ramp-and-hold temporal profile. Candidate source neural signals in the brain might exhibit a ramp-and-hold (tonic) pattern, suggesting a ramp-and-hold-like underlying effect on an as-yet-unknown peripheral mechanism, or could derive from other known temporal profiles including bursts of activity time-locked to saccades. B. Working conceptual model. The brain causes the eyes to move by sending a command to the eye muscles. Each eye movement shifts the location of visual stimuli on the retinal surface. A copy, possibly a highly transformed one, of this eye movement command is sent to the ear, altering ear mechanics in some unknown way. When a sound occurs, the ascending signal to the brain will depend on the combination of its location in head-centered space (based on the physical values of binaural timing and loudness differences and spectral cues) and aspects of recent eye movements and fixation position. This hybrid signal could then be read-out by the brain.

Available eye movement control signals in the oculomotor system include those that follow this ramp-and-hold temporal profile, or tonic activity that is proportional to eye position throughout periods of both movement and fixation. In addition to such tonic signals, oculomotor areas also contain neurons that exhibit burst patterns, or elevated discharge in association with the saccade itself, as well as combinations of burst and tonic patterns [for reviews, see 40, 41]. It remains to be seen which of these signals or signal combinations might be sent to the auditory periphery and where they might come from. The paramedian pontine reticular formation (PPRF) is a strong candidate, having been implicated in providing corollary discharge signals of eye movements in visual experiments [42] [see also 43], and containing each of these basic temporal signal profiles [40, 41]. Regardless of the source and nature of the descending corollary discharge signal, the oscillations observed here should be thought of as possibly constituting a biomarker for an underlying, currently unknown, mechanism, rather than necessarily the effect itself.

Despite these critical unknowns, it is useful to articulate a working conceptual model of how EMREOs might facilitate visual and auditory integration (Figure 9b). The general notion is that, by sending a copy of each eye movement command to the motor elements of the auditory periphery, the brain keeps the ear informed about the current orientation of the eyes. If, as noted above, these descending oculomotor signals cause a ramp-to-step change in the state of tension time-locked to the eye movement and lasting for the duration of each fixation period, they would effectively change the transduction mechanism in an eye position/movement dependent fashion. In turn, these changes could affect the latency, gain, or frequency-filtering properties of the response to sound. The signal sent to the brain in response to an incoming sound would ultimately reflect a mixture of the physical cues related to the location of the sound itself - the ITD/ILD/spectral cues - and eye position/movement information.

Most neurophysiological studies report signals consistent with a hybrid code in which information about sound location is blended in a complex fashion with information about eye position and movement, both within and across neurons [5, 9, 10, 17, 18, 20, 33]. Computational modeling confirms that, in principle, these complex signals can be “read out” to produce a signal of sound location with respect to the eyes [9]. However, substantive differences do remain between the observations here and such neural studies, chiefly in that the neural investigations have focused primarily on periods of steady fixation. A more complete characterization of neural signals time-locked to saccades is therefore needed [7, 44].

Note that this working model differs from a spatial attention mechanism in which the brain might direct the ears to listen selectively to a particular location in space. Rather, the response to sounds from any location, would be impacted by this eye movement/position dependence in a consistent fashion across all sound locations. Such a system may well work in concert with top-down attention, which has previously been shown to impact outer hair cells even when participants are required to fixate and not make eye movements [45–51].

One question that has been explored previously concerns whether sound localization is accurate or inaccurate when a brief sound is presented at the time of an eye movement. Boucher et al [2] reported that perisaccadic sound localization is quite accurate, which suggests that EMREOs (or their underlying mechanism) do not impair perception. This is an important insight because given the frequency and duration of eye movements - about 3/sec – and with each associated EMREO signal lasting 100 ms or longer, it would be highly problematic if sounds could not be accurately detected or localized when they occur in these time windows.

Finally, the findings from our present study argue against another theory: that EMREOs might actually constitute the sounds of the eyeballs moving in their sockets, transmitted to the ear canal via tissue/bone conduction and picked up with the microphone. Several aspects of these results are inconsistent with this hypothesis. The most obvious point is that the EMREO signal persists for many 10’s of ms after the saccade has stopped. A second point is that the EMREO signal in a given ear depends critically on the direction of the associated eye movement, but presumably any summed sound of the left and right eyes moving in conjugate fashion is symmetric: a 10 degree saccade involving both eye balls moving to the left and a 10 degree saccade with both moving to the right should make the same aggregated eye-in-orbits sounds, but they produce EMREO signals that are opposite in phase in each ear canal.

Overall, how brain-controlled mechanisms adjust the signaling properties of peripheral sensory structures is critical for understanding sensory processing as a whole. Auditory signals are known to adjust the sensitivity of the visual system via sound-triggered pupil dilation [52], indicating that communication between these two senses is likely to be a two-way street. The functional impact of such communication at low-level stages is yet to be fully explored and may have implications for how individuals compensate when the information from one sensory system is inadequate, either due to natural situations such as noisy sound environments or occluded visual ones, or due to physiological impairments in one or more sensory systems.

## Methods

### General

Healthy human subjects that were 18 years of age or older with no known hearing deficits or visual impairments beyond corrected vision were recruited from the surrounding campus community (N=10; 5 female, 5 male). If subjects were unable to perform the saccade task without vision correction, they were excluded from the study. All study procedures involving subjects were approved by the Duke Institutional Review Board, and all subjects received monetary compensation for their participation in the study.

Acoustic signals in both ear canals were measured simultaneously with Etymotic ER10B+ microphone systems coupled with ER2 earphones for sound stimulus presentation (Etymotic Research, Elk Grove Village, IL). A low-latency audio interface (Focusrite Scarlett 2i2, Focusrite Audio Engineering Ltd., High Wycombe, UK) was used for audio capture and playback through the Etymotic hardware at a sampling rate of 48kHz. Eye tracking was performed with an Eyelink 1000 system sampling at 1000Hz. Stimulus presentation and data acquisition were controlled through custom scripts and elements of The Psychophysics Toolbox in MATLAB, with visual stimuli presented on a large LED monitor.

In all experiments, eye position and microphone data were recorded while subjects performed silent, visually-guided saccade tasks. Experimental sessions were carried out in a darkened, acoustically isolated chamber (IAC) made anechoic with the addition of acoustic damping wall tiles (Sonex). Subjects were seated and used an adjustable chin rest, positioned 70 cm from the screen, to minimize head movement. Experimental sessions were subdivided into multiple runs, approximately 5 minutes each. This provided subjects with the opportunity to take a brief break from the experiment if needed to maintain alertness or to address discomfort from maintaining their posture. Each run typically consisted of approximately 120 trials and saccade targets were presented in pseudorandom order.

Before each experimental session, the eye-tracking system was calibrated using the calibration routine provided with the Eyelink system to register raw eye-tracking data to gaze locations on the stimulus presentation screen. If the subject requested an adjustment to the chin rest or left the recording chamber for a break, the eye-tracking calibration was repeated. Before each run, the microphone system was calibrated to ensure that each microphone had a frequency response that was similar to the pre-recorded frequency response of the microphones when placed in a volume that approximated the geometry of the human ear canal (see supplementary methods). It consisted of a 3ml syringe cut to accept the Etymotic earpieces. The syringe stopper was pulled to 1.25 cm^3^ to approximate the volume of the human ear canal. A small amount of gauze (.25cm^3^) was added to the volume to emulate the attenuation caused by the soft tissue of the ear canal. The calibration routine played tones from 10 to 1600Hz, at a constant system output amplitude. As the purpose of this calibration was to compare microphone function in a control volume with that in an earpiece just placed in a subject, the weighting of the tones was not specifically calibrated. If the input-output results of the same tone sequences were consistent between ears and matched the overall shape of the syringe calibration curves, microphone placement was considered successful. No sounds were delivered during periods of experimental data collection.

### Task descriptions

All visually-guided saccade tasks followed the same stimulus timing sequence: initial fixation points (referred to as fixation) were displayed on screen for 750ms and then removed as the saccade targets (referred to as target) were presented for 750ms (Figure 1a). Fixation and target locations were indicated by green dots that were 1.5cm in diameter. Subjects were instructed to fixate on the initial fixation locations until targets were presented on the screen, then to saccade to the targets and fixate on the targets until they changed from green to red for the last 100ms of the target presentation at which point the targets were removed from screen (the color change was intended to help subjects maintain consistent fixation periods). Inter-trial-intervals were jittered 350±150ms. This was done to minimize the potential impact of non-saccade related noise signals that may be periodic (i.e. heartbeat, external acoustic and electromagnetic sources).

In the three origin horizontal slice task (Figure 1B), participants performed saccades to multiple targets from three different initial eye positions at −12°, 0°, and +12° horizontally and at 0° of elevation from central fixation. Nine saccade targets were located at 9° elevation and their horizontal locations ranged from −24° to +24° in 6° degree increments

In the criss-cross task (Figure 1C), participants performed saccades to targets along the vertical and horizontal axes from central fixation. Vertical targets ranged from −15° to +15° in 3° increments and horizontal targets ranged from −18° to +18° in 3° increments.

In the (Figure 1D), participants made saccades to 24 distinct targets of varying vertical and horizontal placement combinations from a central fixation. Horizontal location components ranged from −18° to +18° in 9° increments and vertical location components ranged from −12° to +12° in 6° increments.

### Preprocessing analysis

#### Saccade-microphone synchronization

Microphone data was synchronized to two time points in the eye position data: the onset and offset of the targeting saccade. These points in time were defined based on the third derivative of eye position, or jerk. The first peak in the jerk represents the moment when the change in the eye acceleration is greatest, and the second peak represents the moment when the change in the eye deceleration is greatest. Prior to each differentiation, a lowpass discrete filter with a 7ms window was used to smooth the data and reduce the effects of noise and quantization error. This filter was normalized, such that its output to a constant series of values equaled those values.

### Trial exclusion criteria

Trials were excluded based on saccade performance and the quality of recordings for both microphone and eye tracking: 1) if subjects made a sequence of two or more saccades to achieve the target; 2) if the eye tracking signal dropped out during the trial (e.g. due to blinks); 3) if the eye movement was slow and drifting, rather than a saccade; 4) if the saccade curved by more than 4.5° (subtended angle); or 5) subjects failed to maintain 200ms of fixation before and after the saccade. If eye tracking dropped samples that prevented the calculation of the saccade onset and offset times, the trial was also excluded. On average these saccade-related exclusion criteria resulted in the exclusion of about 12% of the trials.

Prior to any further analysis, microphone data was downsampled from 48 kHz to 2 kHz sampling rate to reduce processing time given that the previously observed eye-movement related signals of interest are well below 100 Hz [30]. Exclusion based on noise in the microphone recordings was minimal. Within each block of trials, the mean and standard deviation of the RMS values for each trial was calculated. Individual trials were excluded if the microphone signal on that trial contained any individual values that were more than 10 standard deviations away from that mean. This typically resulted in the exclusion of < ~2% of the trials, after application of the eye position screen described above.

### Z scoring

To facilitate comparison across subjects, sessions, and experiments, all microphone data reported in this study was z-scored within blocks and prior to the application of the exclusion criteria described above. The mean and standard deviation of the microphone values in a window −150 to −120 ms prior to saccade onset was used as the normalization baseline period. When saccade onset- and offset-aligned data were compared with each other (e.g. Figure 2), the same onset-aligned normalization window was used for both.

### Regression Analyses

Regression was used to assess how EMREOs vary with both eye position and eye movement. The microphone signal at each moment in time *Mic(t)* was fit as follows:

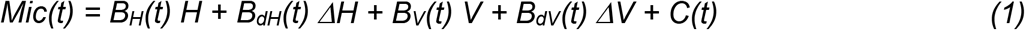

where H and V correspond to the initial horizontal and vertical eye position and *ΔH* and *ΔV* correspond to the respective changes in position associated with that trial. The slope coefficients *B_H_, B_dh_, B_v_, B_dv_* are time-varying and reflect the dependence of the microphone signal on the respective eye movement parameters. The term *C(t)* contributes a time-varying “constant” independent of eye movement metrics, and can be thought of as the best fitting average oscillation across all eye positions and displacements.

The term *C(t)* was included for all regressions, but other parameters were omitted when not relevant. For the grid task and criss-cross tasks, the model used vertical and horizontal saccade displacement (*B_dH_(t) ΔH, B_dv_(t) ΔV)* as regression variables but not *B_H_(t) H* or *B_v_(t) V* as initial position did not vary for those tasks. For the three-origin task, the model included *B_H_(t) H* and *B_dH_(t) ΔH* but not the vertical parameters because those did not vary for this task. However, note that this task did involve a vertical change in position, so the *C(t*) term for this task would tend to incorporate the contributions related to that fixed vertical component.

The regressions for all experiments were carried out twice: once with data synchronized to saccade onset and once with data synchronized to saccade offset. The analysis produced values for the intercept and variable weights (or slopes), their 95% confidence intervals, R^2^, and p-value for each time point.

For the target readout analysis described in Figures 7 and 8, the horizontal and vertical positions of the targets, rather than the associated eye movements, were used.

## Acknowledgments

We are grateful to Dr. Matthew Cooper, Dr. Kurtis Gruters, Dr. David Kaylie, Dr. Jeff Mohl, Meredith Schmehl, Dr. David Smith, Chloe Weiser and Shawn Willett for discussions and assistance concerning this project. This work was supported by NIH (NIDCD) grant DC017532 to JMG.

